# Zinc Oxide Nanoparticles (ZnONPs) as Nanofertilizer: Improvement on Seed Yield and Antioxidant Defense System in Soil Grown Soybean (*Glycine max* cv. Kowsar)

**DOI:** 10.1101/2020.04.13.039644

**Authors:** Elham Yusefi-Tanha, Sina Fallah, Ali Rostamnejadi, Lok Raj Pokhrel

## Abstract

Herein, we investigated potential phytotoxicity of zinc oxide nanoparticles (ZnONPs) on seed yield, focusing on particle size-, morphology-, and concentration-dependent responses of multiple antioxidant defense biomarkers, in soil-grown soybean (*Glycine max* cv. Kowsar) during its lifecycle. To this end, we synthesized three types of morphologically unique ZnONPs (spherical/ 38nm, floral-like/ 59nm, and rod-like/ >500nm); all with high purity, triclinic crystal structure and negative surface charge; and compared the toxicity with Zn^2+^ ions. Each pot received two seeds, placed in soil inoculated with N-fixing bacteria (*Rhizobium japonicum*) and grown outdoor for 120 days. Our findings demonstrated a significant particle size-, morphology-, and concentration-dependent influence of ZnONPs on seed yield, lipid peroxidation, and various antioxidant biomarkers in soybean. Our spherical 38nm ZnONPs were the most protective compared to the floral-like 59nm ZnONPs, rod-like >500nm ZnONPs, and Zn^2+^ ions, particularly up to 160 mg/kg. However, at the highest concentration of 400 mg/kg, spherical 38nm ZnONPs elicited the highest oxidative stress responses (H_2_O_2_ synthesis, MDA, SOD, CAT, POX) in soybean compared to the other two morphologically different ZnONPs tested. The concentration-response curves for the three types of ZnONPs and Zn^2+^ ions were nonlinear (nonmonotonous) for all the endpoints evaluated. The results also suggest differential nano-specific toxicity of ZnONPs compared to ionic Zn^2+^ toxicity in soybean. Our higher NOAEL value of 160 mg/kg indicates the potential for ZnONPs to be used as a nanofertilizer for crops grown in Zn-deficient soils to improve crop yield, food quality and address malnutrition, globally.

**Highlights:**

1. Particle size-, morphology-, and concentration-dependent effects of ZnONPs tested.
2. All Zn compounds (ZnONPs, Zn^2+^) promoted seed yield up to 160 mg/kg.
3. Spherical 38nm ZnONPs elicited the least oxidative stress, except at 400 mg/kg.
4. Concentration-response curves for all Zn compounds were non-linear.
5. ZnONPs may serve as nanofertilizer for enriching Zn-deficit soil with Zn.

TOC Art

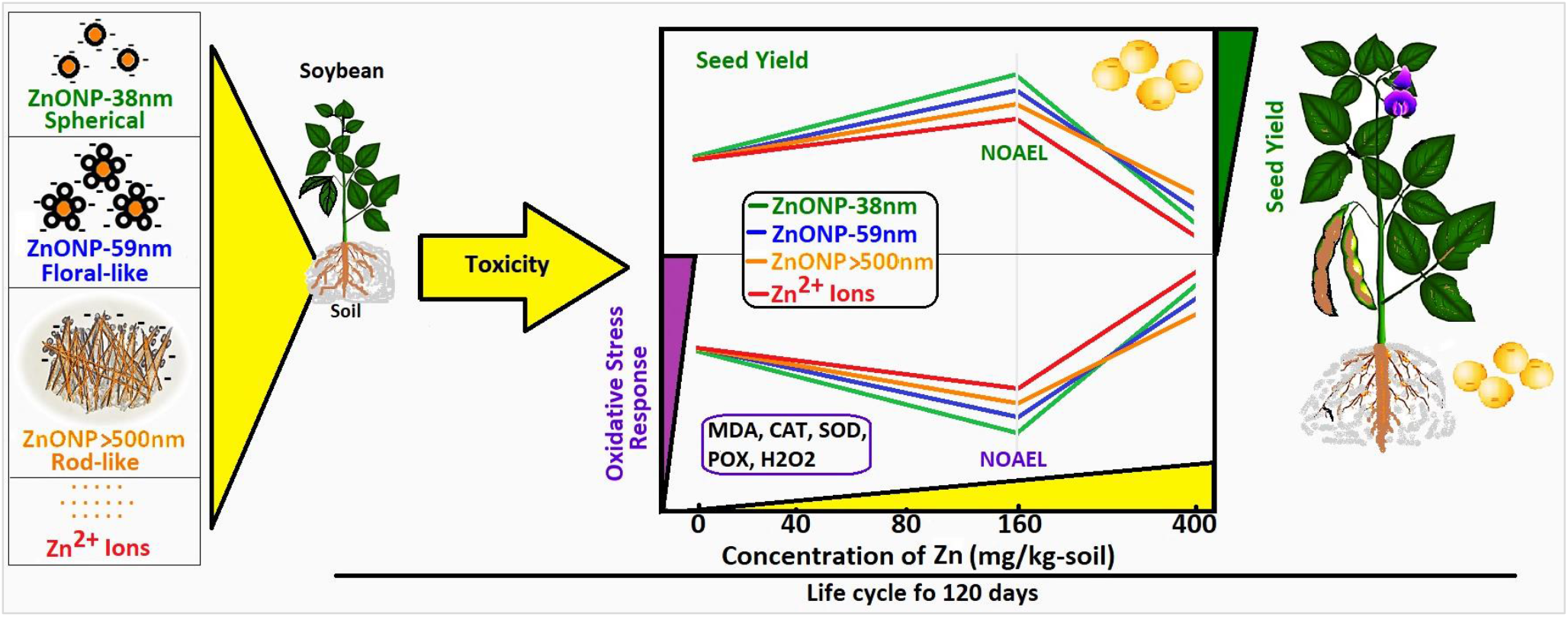

## Introduction

Nanoscience and nanotechnology imply the study of engineered nanostructures with at least one dimension below 100 nm with potential for application in a variety of sectors, including in agriculture, therapeutics, diagnostics, engineering, food industry and safety, environmental remediation, and energy infrastructure, among others (National Nanotechnology Initiative, 2005; Du et al., 2019; Malea et al., 2019). Production of engineered nanoparticles (ENPs) for the said applications is exponentially growing primarily due to unique nano-specific physicochemical properties such as high surface area to volume ratio, greater reactivity, high electrical conductivity, high mechanical strength, and antimicrobial properties (Hou et al., 2018; Pokhrel et al., 2012; Rajput et al., 2018).

Growing interests in the use of ENPs in food safety and agriculture can also be attributed to their unique size and surface-related behavior (Nuruzzaman et al., 2016; Dimkpa et al., 2019). ENPs may find use in agriculture in the form of nanofertilizers, growth regulators, or as nanopesticides to address plant and zoonotic diseases. Further, ENPs may serve as sensors in early detection of pathogens and/or degradation of pesticide residue due to higher reactivity (Wang et al., 2016b; Xiong et al., 2017). ZnONPs with unique physiochemical properties may serve as a novel fertilizer for the improvement in crop yield and food quality (Dimkpa and Bindraban, 2018; Hou et al., 2018; White and Gardea-Torresdey, 2018).

Plants are primary producers in virtually every ecosystem, upon which consumers rely (Ma et al., 2010). As such, plants can also uptake and transport various toxicants including ENMs from the biosolid, soil and irrigation water into the edible parts (root, leaf, flower, fruit, seed) (Andreotti et al., 2015; Ogunkunle et al., 2018; Yusefi-Tanha et al., 2020) and serve as a source of exposure to humans and other organisms that feed upon plants. Eventually, toxicants may transport through different trophic levels in the food chain (Zhu et al., 2008; Ghasemi Siani et al., 2017). ENPs may interact with the root surfaces while in the soil or hydroponic media, from where ENPs may traverse through apoplastic and symplastic pathways within the plant cells and thence move up to shoot via the xylem vessel (Deng et al., 2014; Lin and Xing, 2008; Lin et al., 2009; Schwab et al., 2016). It is beginning to be understood that ENPs toxicity on plant growth, metabolism, defense system, and yield may depend on the route of exposure, media composition, dose, particle morphology (size, shape), particle composition, and surface chemistry (Cao et al., 2017; Deng et al., 2017; Pagano et al., 2017; Pokhrel et al., 2012; del Real et al., 2017), including the chemical makeup of the subcellular sites and plant bioassays (Dietz and Herth, 2011; Lin and Xing, 2008; Nuruzzaman et al., 2016; Kranjc et al., 2018; Xiong et al., 2017). Improving knowledge on potential factors influencing phytotoxicity, phytotoxicity mechanisms and mitigation measures will support future risk assessment of ENPs.

Several studies have reported positive effects of nanoparticles on plant germination, growth and performance. For example, increased seed germination and growth of seedling, improved photosynthetic efficiency, biomass, and total protein, sugar, nitrogen, and micronutrients were observed in several crop plants; e.g., *Vigna radiata* and *Cicer arietinum* (Mahajan et al., 2011), *Spinacia oleracea* (Srivastava et al., 2014), *Cucumis sativus* (Moghaddasi et al., 2017), *Solanum lycopersicum* (Faizan et al., 2018), and *Triticum aestivum* (Zhang et al., 2018). *C. sativus* grown in gel chamber showed increased shoot and root biomass with ZnONPs (1 mg/L), and increased shoot length with ZnONPs compared to bulk ZnO (Moghaddasi et al., 2017). When ZnONPs and their derivatives, including ZnCl_2_, are available in soil in excess, potential toxicity may result in plants (Liu et al., 2015; Mukherjee et al., 2014a; Wang et al., 2016a), including inhibitory effects on seed germination, growth, photosynthesis, physiological and biochemical traits, yield characteristics, and nutritional quality (Du et al., 2017; Zuverza-Mena et al., 2017).

Previous studies have reported potential oxidative stress upon exposure to different ENPs in plants (Mukherjee et al., 2016). Oxidative stress ensues when the equilibrium between reactive oxygen species (ROS) production and defense system in plants is impaired. ROS, which are produced as the by-product of light reaction in chloroplasts during photosynthesis, are implicated in the toxicity elicited by ENPs (Ma et al., 2015) as excess ROS may lead to DNA damage, oxidation of protein, peroxidation of lipid, membrane damage, electrolyte leakage, and eventually, cell death (Demidchik, 2015; Wang et al., 2018). To maintain homeostasis, plants have evolved various strategies that enable protection of cellular and sub-cellular components from potential toxicity of free radicals via deactivating or removing the ROS. These mechanisms consist of both non-enzymatic antioxidants, including the low molecular weight compounds (phenolics, ascorbate, a-tocopherols, glutathione, carotenoids, and proline), and enzymes such as superoxide dismutase (SOD), catalase (CAT), peroxidase (POD), and ascorbate peroxidase (APX), among others (Getnet et al., 2015; Ozyigit et al., 2016). These antioxidants reduce ENP related oxidative injury in plants (Siddiqi and Husen, 2017). MDA is one of the by-products of oxidative stress and ROS that results from the oxidation of unsaturated fatty acids on the cell membrane (Gaschler and Stockwell, 2017).

Recently, we showed particle size (25, 50 and 250 nm)- and concentration-dependent effects of CuONPs on lipid peroxidation, and antioxidant biomarkers including SOD, CAT, POX, and APX, in soybean grown in mycorrhiza-enriched soil for a period of 120 days in the outdoor environment (Yusefi-Tanha et al., 2020). Particularly, the effects of 25nm CuONPs were found to be consistently higher for most of the antioxidant biomarkers tested compared to the two larger size CuONPs (CuONPs 50nm, CuONPs 250nm), or Cu^2+^ ions, treatments. We also showed that the concentration-response curves for 25nm CuONPs and Cu^2+^ ions were linear, unlike for the larger size CuONPs (CuONPs 50nm, CuONPs 250nm) the relationships were nonlinear, for most of the antioxidant biomarkers assessed. The concentration-response curves for seed yield for all types of Cu compounds were linear (R^2^>0.65). Soybean seed yield also showed particle size- and concentration-dependent inhibition with CuONPs, and inhibition of 25nm CuONPs were significantly higher than the two larger size CuONPs or Cu^2+^ ions at all concentrations tested. Our results indicated differential nano-specific toxicity compared to ionic Cu^2+^ toxicity in soybean (Yusefi-Tanha et al., 2020).

Zinc (Zn) is a necessary micronutrient for all plants as it plays a vital role in many physiological activities such as in the biosynthesis of chlorophyll, proteins and enzymes, including in metabolic processes (Singh et al., 2018). For example, Zn is present within the cytosolic and chloroplastic copper/zinc-SOD enzymes that play a critical role against oxidative stress (Yruela, 2015). Zn metalloproteins are known to partake in replication and transcription of DNA, thereby regulating gene expression (Barker and Pilbeam, 2015). In addition, Zn is a structural part of the ribosome and responsible for its structural integrity, and takes part in amino acid synthesis and nitrogen metabolism (Barker and Pilbeam, 2015; Kobraee and Shamsi, 2015). Zn is also found in the structure of CCCH-type Zn finger proteins, that genes encoding them increase the oil content in soybean seed via activating genes related to lipid biosynthesis (Li et al., 2017). In addition, Zn stimulates N2 fixation in legumes, such as French bean (*Phaseolus vulgaris* L.), by promoting nodule number in roots (Hemantaranjan and Garg, 2015). Deficiency of Zn in plants is a matter of global concern (Impa et al. 2013), thereby highlighting the potential of supplementing Zn in the form of nanoparticles as fertilizer (Dimkpa et al., 2015; Du et al., 2019). However, it is noted that at higher concentrations ZnONPs may also cause toxicity in plants (Dimkpa et al., 2015; Liu et al., 2015, wang et al., 2016a).

Soybean, one of the most cultivated and economically important food crops, is rich in protein and oil content and is consumed globally (FAO, 2019; Kanchana et al., 2016). Soil fertilization can affect the nutritional quality of the crop seed such as soybean (Zulfiqar et al., 2019). In this study, we investigated if zinc oxide nanoparticles (ZnONPs) could serve as a novel nanofertilizer, one that could improve crop health and productivity. We tested particle size-, morphology- and concentration-dependent responses on seed yield and antioxidant defense system in soil grown soybean (*Glycine max* cv. Kowsar) during its lifecycle of 120 days. To achieve this goal, we synthesized three types of morphologically different ZnONPs (spherical/ 38nm, floral-like/ 59nm and rod-like/ >500nm); all with high purity, triclinic crystal structure and negative surface charge.

## Material and methods

### Synthesis and characterization of ZnO nanoparticles

ZnONPs with three different morphologies (spherical/ 38nm, floral-like/ 59nm and rod-like/ >500nm) were prepared by ball milling and sol-gel methods. Analytical grade zinc acetate dehydrate (Zn(CH_3_COO)_2_.2H_2_O) and citric acid (C_6_H_8_O_7_) were purchased from Merck and used as received. The details of the synthesis process were previously described by Zandi et al. (2011). Briefly, zinc acetate and citric acid (as powder) were mixed in a molar ratio of 1:1 and ground for 1 h at room temperature. The milled powder was calcinated at 530°C for 10 h to obtain ZnONPs (S_1_). Samples of ZnONPs with larger particle sizes were also prepared by sol gel method. For this purpose, stoichiometric amounts of zinc acetate and citric acid in the molar ratio of 1:1 were dissolved in distilled water. The solution was stirred using a magnetic stirrer at 80 °C when a viscous gel was obtained. The gel was dried at 100 °C to transform into powder, then divided into two batches; each batch was calcinated at 800 °C (S_2_), or 1000 °C (S_3_) in the air atmosphere. Phase formation and crystal structure of the samples were characterized using X-ray diffraction (XRD) in the scanning angle range of 2θ=20-80° using a Philips X’Pert PRO X-ray diffractometer by Cu-Kα X-ray source (λ=1.5406 Å). Field emission-scanning electron microscopy (FE-SEM) was used to determine particle morphology and diameter. Dynamic light scattering (DLS) was used to estimate hydrodynamic diameter (HDD) and zeta ζ) potential (of the ZnONPs synthesized.

### Experimental set up

Our experiments followed a two-way factorial design, which comprised of: ZnCl_2_ (Zn^2+^; positive control) and ZnONPs with three different morphologies (ZnONP; spherical/ 38nm, floral-like/ 59nm, and rod-like/ >500nm) and five different concentrations (0, 40, 80, 160 and 400 mg Zn/kg). To avoid any spatial effects, the treatments followed a randomized complete design (RCD) with three replicates per treatment, for a total of 60 experimental units with two plants each (*n* = 120 plants). All experiments were carried out at Shahrekord University, Iran.

### Soil characterization

Soil was collected from the surface layer (0-30 cm depth). In order to separate wood chips, large clots and rocks, the soil was air-dried and sieved (2 mm mesh). The concentration of background zinc in soil was 0.892 mg/kg. The main physicochemical characteristics of this soil are as follows: classified as silt loam soil (16% sand, 58% silt, and 26% clay), with a pH of 7.44, EC of 0.47 dS/m, 9.24 g/kg of organic carbon, 0.88 g/kg of total N, and 11.7 and 405 mg/kg of available P and K, respectively.

### Soil amendment with zinc compound

For soil amendment, each zinc compound (Zn^2+^ and ZnONP: spherical/ 38nm, floral-like/ 59nm and rod-like/ >500nm) were weighed and suspended in 100 mL of distilled water to achieve desired concentrations (0, 40, 80, 160 and 400 mg Zn/kg soil). ZnONPs and Zn^2+^ ions suspensions were stirred with a magnetic bar for 30 min at 25°C before being added to the soil. Then, different Zn treatments (ZnONP and Zn^2+^) were added to the soil. ZnCl_2_ was added to soil as an aqueous solution (positive control) to determine potential effects of Zn^2+^ to soybean. Untreated soil served as the negative control. Three replicates of each treatment were prepared. ZnONPs and Zn^2+^ ions in aqueous suspensions were hand mixed with 1 kg of soil for 30 mins, repeated mixing thrice with 1 kg soil each time for a total of 4 kg of soil per pot (to enable homogeneous mixing), and equilibrated in outdoor environment for 24 h before planting ensued.

### Crop management through maturity and seed production

Briefly, control and amendment soil samples (4 kg) were placed in PE (polyethylene) bags and put in (20 cm diameter × 20 cm height) PE pots. Each pot had an inner liner of PE mesh (all bags had 50 holes of 5 mm for drainage), and bottom-filled with a layer of washed gravel (500 g), facilitating aeration and drainage. This design enabled the root system to remain within the bag/pot and facilitated easier plant removal from pot at harvest. Soybean seeds (Kowsar cultivar) were obtained from the Seed and Plant Improvement Institute, Karaj, Iran. They were hydrated in distilled water for 24 h to facilitate germination. Two seeds with uniform size were planted after inoculation with soybean symbiotic bacteria (*Rhizobium japonicum*) at 2.5 cm soil depth 24 h after soil amendment. This study was performed in microcosm conditions to better understand the real effects of NPs in the environment. During the full growth period, the pots were irrigated based on field capacity. At each irrigation, a sub-sample of water was measured for Zn concentration by inductively coupled plasma-optical emission spectroscopy (ICP-OES). Upon maturity (120 days post-planting), the plants and seeds were harvested. Seeds were air-dried, and weight recorded.

### Determination of biochemical parameters

Two youngest leaves per pot (one leaf each of two plants per pot) were sampled to determine all biochemical parameters tested (Yusefi-Tanha et al., 2020).

### Measurement of malondialdehyde concentration

The level of lipid peroxidation was determined by measuring the formation of malondialdehyde (MDA) content with thiobarbituric acid (TBA) by using the method of Heath and Packer (1968). Briefly, fresh leaf tissue samples (0.1 g) was homogenized in 1.5 ml of 0.1% trichloroacetic acid (TCA). The resultant homogenate was centrifuged at 10,000×g for 10 min, and 1 mL of the supernatant was added to 2 ml of 20% TCA containing 0.5% TBA. The extract was heated in a water bath at 95°C for 30 min and then quickly cooled in an ice bath. After that, centrifuged at 10,000×g for 10 min. The absorbance of the supernatant was read at 532 and 600 nm against a blank. MDA concentration was expressed in terms of nmol/g FW (by using extinction coefficient of 155 mM^-1^ cm^-1^) (Narwal et al., 2009).

### Estimation of hydrogen peroxide (H_2_O_2_)

The hydrogen peroxide produced in leaf of plants was measured following the method previously described by Nag et al. (2000). Fresh leaf tissue (1 g) was powdered using liquid nitrogen and was homogenized in 12 mL cold acetone. Then, homogenate was filtered through Whatman filter paper. The mixture was diluted using 4 mL reagent of titanium 16%, and 0.2 mL ammonium hydroxide 28%. The tissue extract was further centrifuged at 8500 rpm for 5 min at 4°C. The supernatant is isolated and then the precipitate was washed twice with 5 mL acetone. Finally, added 2 mL of sulfuric acid (1M), and absorption was measured at 410 nm. The hydrogen peroxide content was calculated using the standard curve prepared in the similar way and expressed as nmol/g FW.

### Estimation of superoxide dismutase (SOD)

The tissue samples (1 g of fresh leaf tissue) were frozen in liquid nitrogen and homogenized in 10 mL of 0.1 M potassium phosphate buffer (pH 7.5). The tissue extract was further centrifuged at 20,000 rpm for 30 min at 4°C. The supernatant was collected, separated into several aliquots and stored at −20°C for determination of antioxidant enzyme activities. The SOD activity was determined by measuring inhibition of the photochemical reduction of nitroblue tetrazolium (NBT) (Narwal et al., 2009). 1.95 mL of 0.1 M potassium phosphate buffer (pH 7.5), 250 μL of 150 mM methionine, 250 μL of 1.2 mM Na_2_EDTA, 250 μL of 24 μM riboflavin, 250 μL of 840 μM NBT, and 50 μL of plant extract were prepared. The reaction was initiated by light illumination and the absorbance was read at 560 nm. One unit of SOD activity is defined as the amount of enzyme which causes 50% inhibition of oxidation reactions per milligram of protein in extract.

### Estimation of catalase (CAT) activity

The CAT activity was determined according to Narwal et al. (2009) by measuring decrement in absorbance at 240 nm following the decomposition of hydrogen peroxide (H_2_O_2_). The reaction mixture consisted of 50 μL of supernatant, 1.95 ml of 0.1 M potassium phosphate buffer (pH 7.0) and 100 μL of 264 mM H_2_O_2_. The decrease in absorption was detected for a period of 100 second at 5 second intervals at room temperature (25°C). One unit of CAT activity corresponds to 1 mMol of H_2_O_2_ consumed per min per mg of protein using an extinction coefficient of 40 mM^-1^ cm^-1^.

### Estimation of guaiacol peroxidase (POX) activity

The guaiacol peroxidase (POX) activity was estimated following the method previously developed by MacAdam et al. (1992). Briefly, 50 μL of plant extract was added to 1.35 mL 0.1 M potassium phosphate buffer (pH 6.0), 100 μL 45 mM guaiacole, and 500 μL 44 mM hydrogen peroxide. Then, we measured kinetic of changes in absorbance at 470 nm at 10 second interval for 300 seconds at 25°C using UV-Vis spectrophotometer. One unit of POX activity corresponds to 1 mMol tetraguaiacol consumed per min. per mg of protein using an extinction coefficient of 26.6 mM^-1^ cm^-1^.

### Estimation of ascorbate peroxidase (APX) activity

The APX activity was measured by monitoring the rate of ascorbate oxidation with H_2_O_2_, following the method developed by Narwal et al. (2009). The decrease in ascorbic acid, starting from a mixture of 2.4 mL of 0.1 M potassium phosphate buffer (pH 7.0), 250 μL of 1.2 mM Na2EDTA, 50 μL of 35 mM H_2_O_2_, 100 μL of 15 mM ascorbic acid, and 200 μL of supernatant was measured at 290 nm over a period of 500 s at 10 s interval at room temperature (25°C). The activity was calculated using an extinction coefficient of 2.8 mM^-1^ cm^-1^. One unit of APX was defined as 1 mMol of ascorbate oxidized per min. per mg of protein.

### Estimation of scavenging of H_2_O_2_

The scavenging of H_2_O_2_ was measured following the method by Benkebila (2005), which measures decrement in UV light absorbance at 230 nm due to H_2_O_2_ scavenging activity of tested substances/extracts. Each sample was prepared by adding 1 ml of the plant extract and 600 μL of fresh prepared 2 mM H_2_O_2_ in phosphate buffer saline (PBS). In another tube a control sample was prepared by adding 1 mL of PBS and 600 μL of fresh prepared 2 mM H_2_O_2_ in PBS. Then, the mixtures were left for 10-15 min. at room temperature. Then absorbance was measured at 230nm.

### Statistical analysis

Data were analyzed with two-way analysis of variance (ANOVA) to examine the interaction between the factors using the least significant difference test using SAS (SAS Inc., ver. 9.4). The differences were considered significant at *p*<0.05 and the means were separated using a Fisher’s protected LSD test. The results are presented as the mean ± standard deviation (SD). To determine if the concentration-response curves were linear (monotonic) or nonlinear (nonmonotonic), we coupled visual inspection of the curves with a simple decision rule: if the coefficient of determination (R-squared) value for the linear regression line is 65% or higher the concentration-response curves were deemed linear, suggesting that the plant response changes linearly with the concentration applied following the relationship: y = ax + b; where y denotes dependent variable, x denotes independent variable, and a and b are model parameters. The computed R-squared values are presented in Supplemental Information (SI) **Table S1**.

## Results

### Nanoparticle characterization

The analysis of the XRD patterns show that the samples are in triclinic crystalline form with space group p63mc, without any noticeable trace of impurities. The XRD patterns of the powder samples are shown in **Fig. 1.** The morphology and average particle diameter of the ZnONPs were measured by field emission scanning electron microscopy (FE-SEM). Representative FE-SEM micrographs of the ZnONPs samples and their particle size distributions (PSD) fitted to a lognormal distribution function (as shown in equation 1 below; Rostamnejadi et al., 2017) are presented in **Fig. 1**:

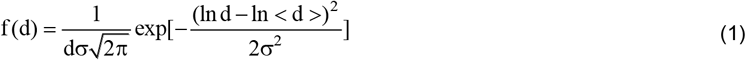

**Fig. 1.**
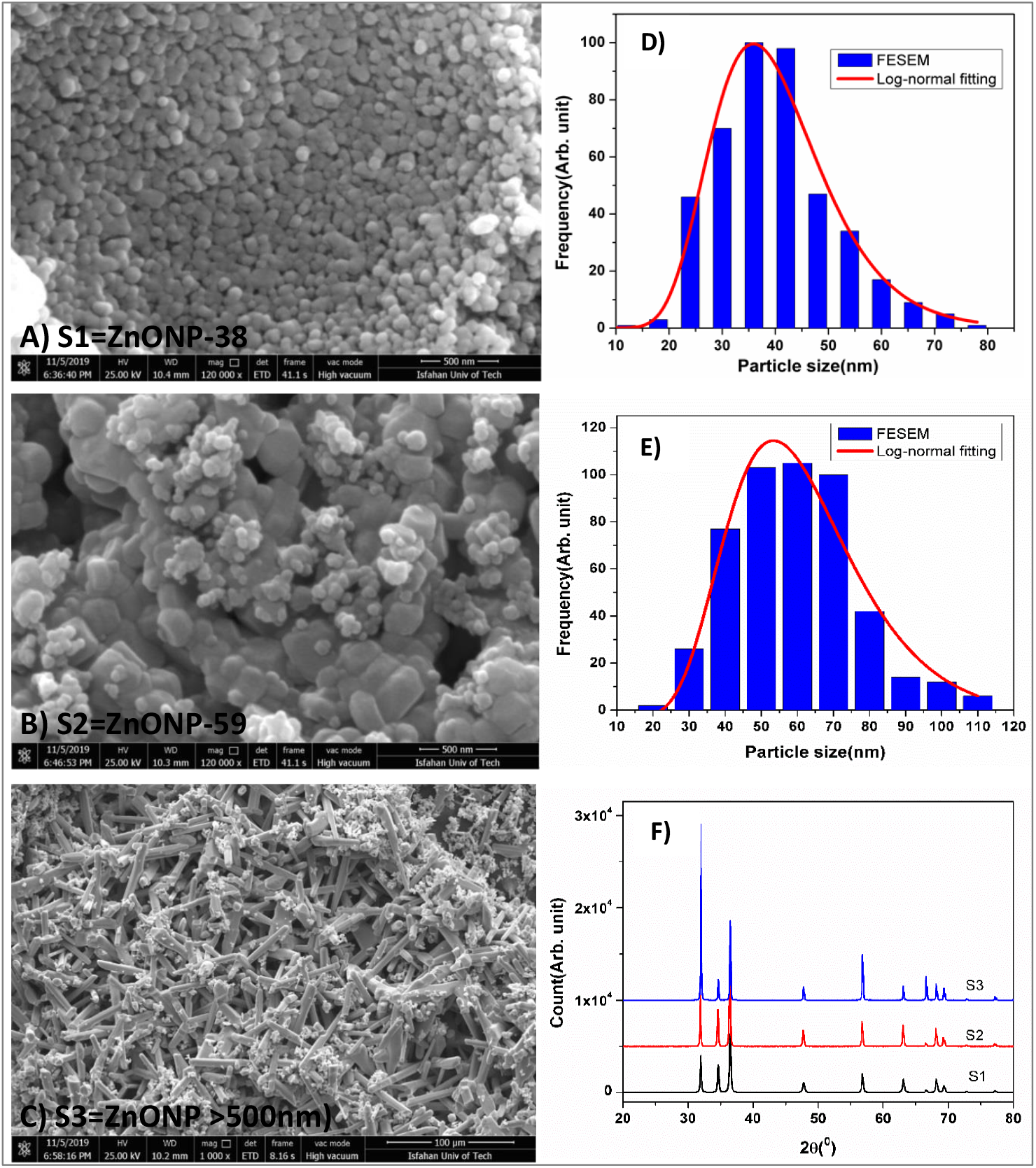
FE-SEM micrographs of the three morphologically distinct ZnONPs: A) S1 (spherical/ 38nm diameter); B) S2 (floral-like/ 59nm diameter); and C) S3 (rod-like/ >500nm diameter and length >1μm). Particle size distribution (PSD) of sample S1 (D) and sample S2 (E). F) XRD patterns of the powder samples of the three morphologically distinct ZnONPs: S1, S2, and S3.

In this relation <d> is the mean particle size and σ is the standard deviation. Our analysis of FE-SEM micrographs of S1 samples showed particles with nearly spherical shape with an average particle diameter of 38 nm but with dense packing appeared as potential aggregation might have occurred. S2 samples presented floral-like morphology with particle diameter of 59 nm, on average. S3 samples showed spherical particles with average diameter larger than 500 nm atop microrods of about 1μm in diameter and 10-50 μm in length. All ZnONPs samples (S1-S3) were highly negatively charged with over 44.0 mV zeta potential but had variable hydrodynamic diameters (HDDs) (**Table 1**), a result consistent with the FE-SEM analyses.

**Table 1.** Dynamic light scattering (DLS) measurement of hydrodynamic diameter (HDD) and zeta potential values for: (S_1_) spherical ZnONP-38nm, (S2) floral-like ZnONP-59nm, and (S3) rod-like ZnONP >500nm.

| Sample | Zeta Potential (mV) | HDD (nm) |
| --- | --- | --- |
| S1 = spherical ZnONP-38nm | -44.8 | 139 |
| S2 = floral-like ZnONP-59nm | -50.4 | 194 |
| S3 = rod-like ZnONP<br>>500nm | -54.6 | 1059 |

### Impacts on seed yield

Our results show the effects of zinc compound types (Zn_type_), zinc concentration (C), and the interaction of Zn_type_ × C were statistically significant on seed yield in soybean (*p*<0.0001) (**Table 2**). Generally, there was concentration-dependent increase in seed yield for all Zn compounds up to 160 mg/kg, but at the highest concentration tested (400 mg/kg) seed yield significantly declined for all Zn compounds (**Fig. 2**). Seed yield was highest for spherical 38 nm ZnONPs compared to floral-like 59 nm or rod-like >500 nm ZnONPs. Seed yield was typically lowest for Zn^2+^ ions treatments compared to all three types of ZnONPs treatments (**Fig. 2**), suggesting inhibitory effects of Zn^2+^ ions on seed yield compared to ZnONPs tested. Further, highest seed yield occurred at 160 mg/kg for all Zn compound types. Overall, the concentration response curves for all types of Zn compounds were non-linear (**Fig. 2;** see SI **Table S1** for R-squared values), and that the responses were particle size- and morphology-dependent.

**Fig. 2.**
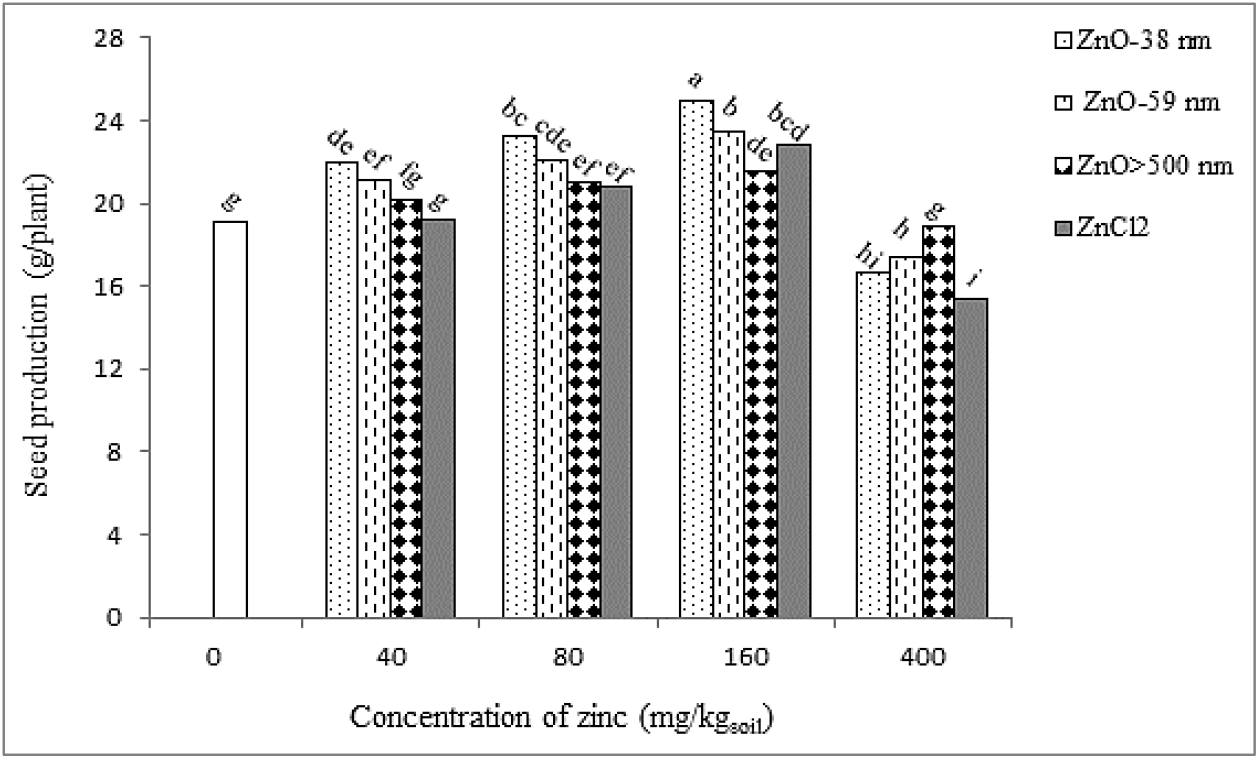
Seed production in soil grown soybean treated with different types of zinc compounds (spherical ZnONP-38nm, floral-like ZnONP-59nm, rod-like ZnONP>500nm, and Zn^2+^ ions) as a function of concentrations (0, 40, 80, 160, 400 mg Zn/kg soil). Error bars represent mean ± SD. Different letters above the bar indicate significant difference at *p*<0.05.

**Table 2.** Analysis of variance (*p-*value) for seed production and biochemical parameters of soybean grown in soil treated with different concentration of zinc compound types.

| Source of variation | Seed production | H<sub>2</sub>O<sub>2</sub> production | H<sub>2</sub>O<sub>2</sub> scavenging | Malondialdehyde (MDA) |
| --- | --- | --- | --- | --- |
| Zinc compound type (Zn<sub>type</sub>) | <.0001 | <.0001 | <.0001 | <.0001 |
| Concentration (C) | <.0001 | <.0001 | <.0001 | <.0001 |
| Zn<sub>type</sub> × C | <.0001 | <.0001 | <.0001 | <.0001 |

### Impacts on hydrogen peroxide (H_2_O_2_) production

Our ANOVA showed the effects of Zn_type_, C, and Zn_type_ × C on leaf H_2_O_2_ production in soybean were significant (*p*<0.0001) (**Table 2**). We observed particle size-, morphology- and concentration-dependent influence of ZnONPs on H_2_O_2_ production in soybean leaf (**Fig. 3**). With increasing concentrations, H_2_O_2_ production significantly decreased up to 160 mg/kg for all Zn compounds; whereas at 400 mg/kg, H_2_O_2_ production increased significantly for all Zn compounds, except for rod-like >500 nm ZnONPs treatment that showed no significant change compared to control (**Fig. 3**). Overall, these results suggest that all Zn compounds were protective to soybean up to 160 mg/kg, and that the responses were non-linear (**Fig. 3;** see **SI Table S1** for R-squared values).

**Fig. 3.**
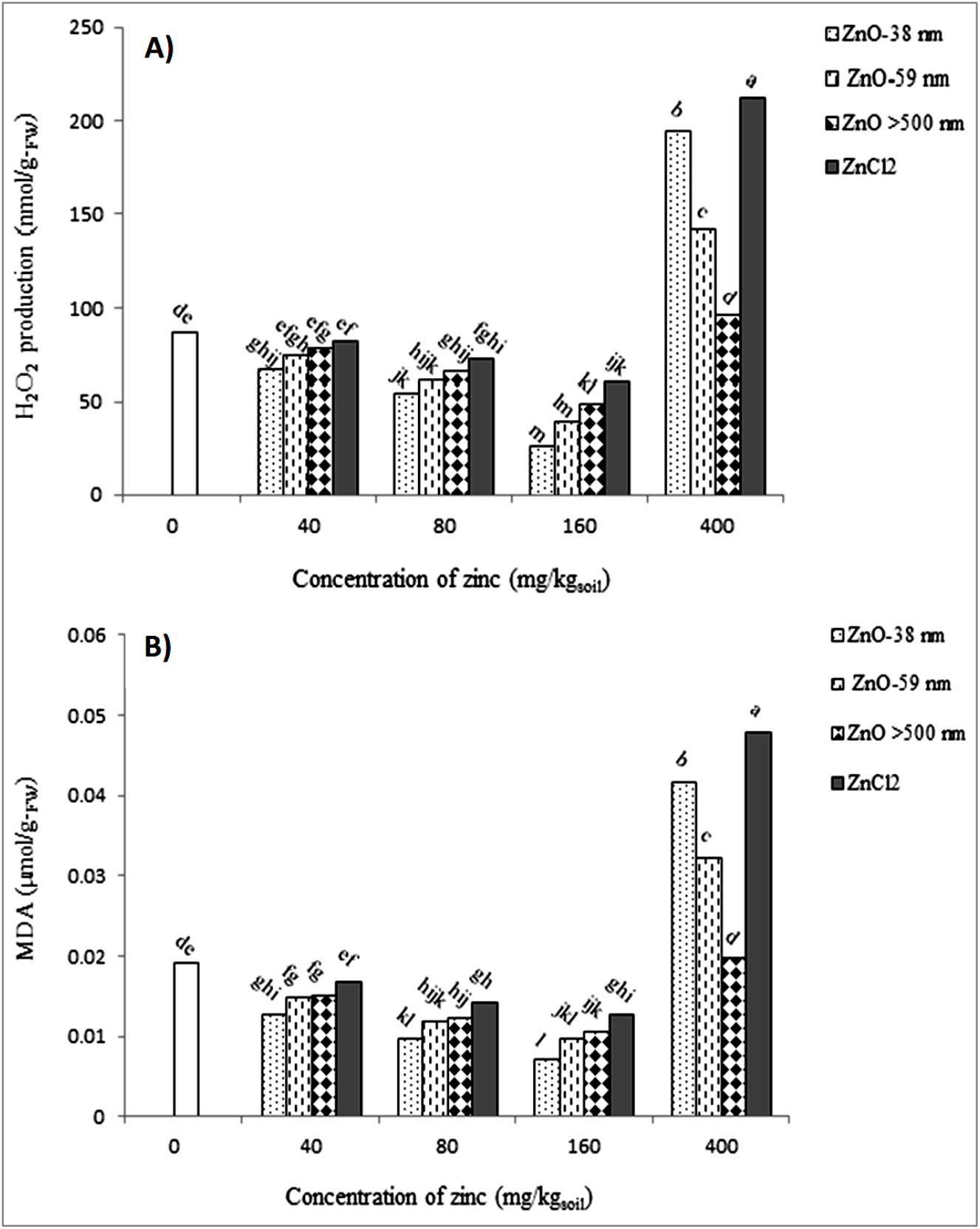
Changes in leaf hydrogen peroxide (H_2_O_2_) production (**A**), and leaf malondialdehyde (MDA) content (**B**), in soil grown soybean treated with different types of zinc compounds (spherical ZnONP-38nm, floral-like ZnONP-59nm, rod-like ZnONP>500nm, and Zn^2+^ ions) as a function of concentrations (0, 40, 80, 160, 400 mg Zn/kg soil). Error bars represent mean ± SD. Different letters above the bar indicate significant difference at *p*<0.05.

### Impacts on malondialdehyde (MDA) accumulation

An accumulation of the MDA-TBA complex was measured to estimate lipid peroxidation in the leaf tissue. Our results showed that MDA content was affected by Zn_type_ (*p*<0.0001), C (*p*<0.0001), and their interactions (Zn_type_ × C) (*p*<0.0001) (**Table 2**). We observed particle size-, morphology- and concentration-dependent influence of ZnONPs on MDA levels in soybean leaf (**Fig. 3**). With increasing concentrations, MDA content significantly decreased up to 160 mg/kg for all Zn compounds; whereas at 400 mg/kg, MDA content increased significantly for all Zn compounds, except for rod-like >500 nm ZnONPs treatment that showed no significant change compared to control (**Fig. 3B**). As expected, the MDA results mirrored the results of H_2_O_2_ production in leaf suggesting reciprocal relation between H_2_O_2_ production and MDA-TBA accumulation upon lipid peroxidation. Overall, the lipid peroxidation responses were non-linear (**Fig. 3;** see SI **Table S1** for R-squared values).

### Impacts on superoxide dismutase (SOD), catalase (CAT) and guaiacol peroxidase (POX) activities

Activities of multiple antioxidant enzymes (SOD, CAT and POX) in soybean leaf exposed to different Zn compounds are presented in **Fig. 4**. The effects of Zn_type_ (*p*<0.0001), C (*p*<0.0001) and the interaction term (Zn_type_ × C) (*p*<0.0001) were statistically significant on the antioxidant enzymes (SOD, CAT, and POX) activities in soybean leaves. Similar to the H_2_O_2_ production and subsequent MDA accumulation documented above, we observed particle size-, morphology- and concentration-dependent influence of ZnONPs on SOD, CAT and POX activities in soybean leaf (**Fig. 4**). With increasing concentrations, these antioxidant enzyme activities were significantly reduced up to 160 mg/kg for all the Zn compounds; whereas at 400 mg/kg, these enzyme activities increased significantly for all Zn compounds, except for rod-like >500 nm ZnONPs treatment that showed no significant difference in the enzyme activities compared to control (**Fig. 4**). Overall, these results suggest that all three antioxidant enzymes’ activities were consistent with the trend for H_2_O_2_ synthesis and MDA accumulation responses, and that the enzymes’ responses were non-linear (**Fig. 4;** see **SI Table S1** for R-squared values).

**Fig. 4.**
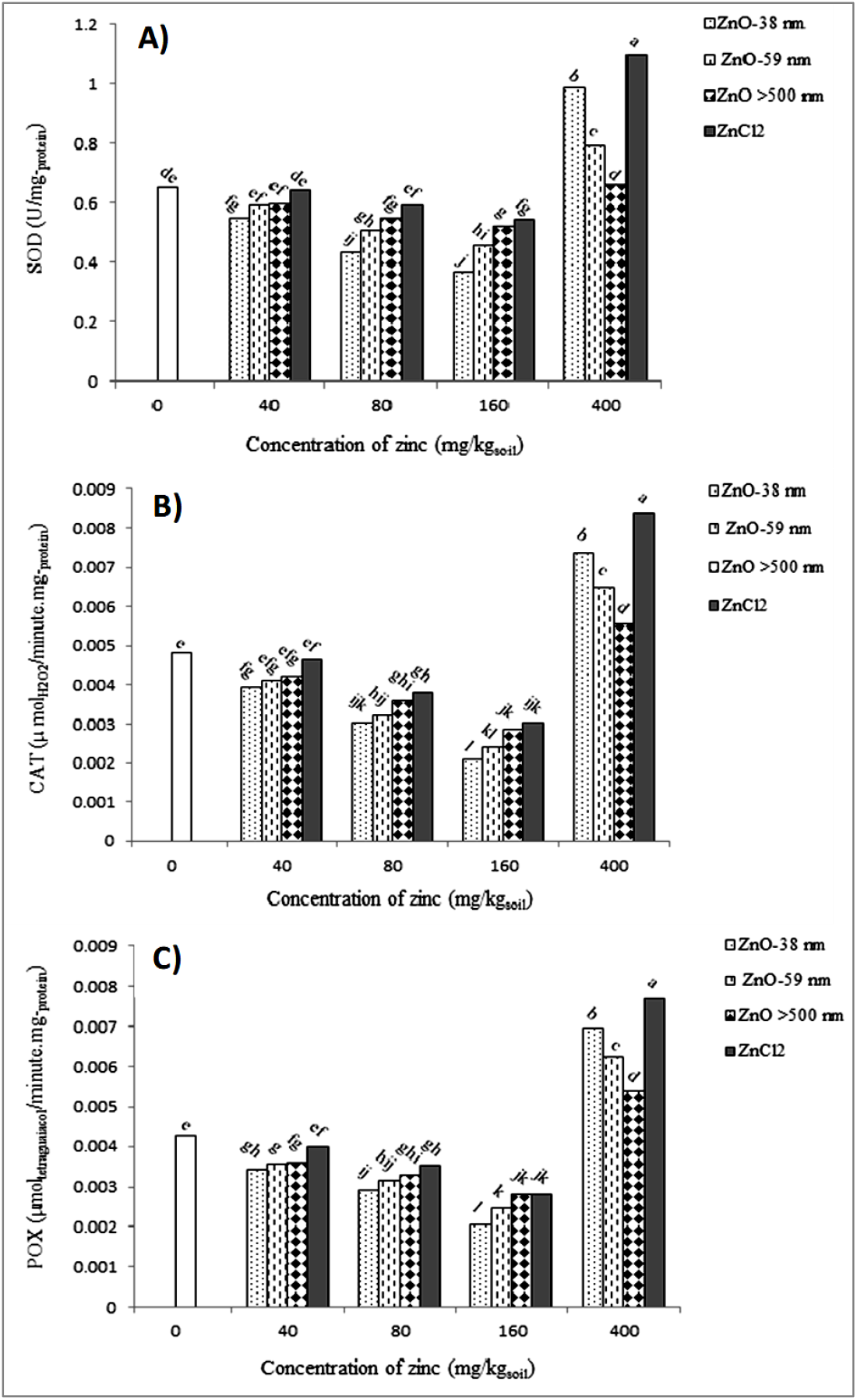
Changes in leaf superoxide dismutase (SOD) (**A**), catalase (CAT) (**B**), and guaiacol peroxidase (POX) (**C**), activities in soil grown soybean treated with different types of zinc compounds (spherical ZnONP-38nm, floral-like ZnONP-59nm, rod-like ZnONP>500nm, and Zn^2+^ ions) as a function of concentrations (0, 40, 80, 160, 400 mg Zn/kg soil). Error bars represent mean ± SD. Different letters above the bar indicate significant difference at *p*<0.05.

### Impacts on ascorbate peroxidase (APX) and H_2_O_2_ scavenging activities

The effects on APX and H_2_O_2_ scavenging activities in soybean leaf exposed to different Zn compounds are depicted in **Fig. 5**. ANOVA showed the effects of Zn concentration (C) and the interaction term (Zn_type_ × C) were statistically significant on APX activity (*p*<0.0001) in soybean leaf. Whereas, the effects of Zn_type_, C and the interaction term (Zn_type_ × C) were all statistically significant on H_2_O_2_ scavenging (*p*<0.0001) in soybean leaf. Overall, the APX responses of soybean upon exposure to different types of Zn compounds were opposite to that of H_2_O_2_ scavenging activities observed.

**Fig. 5.**
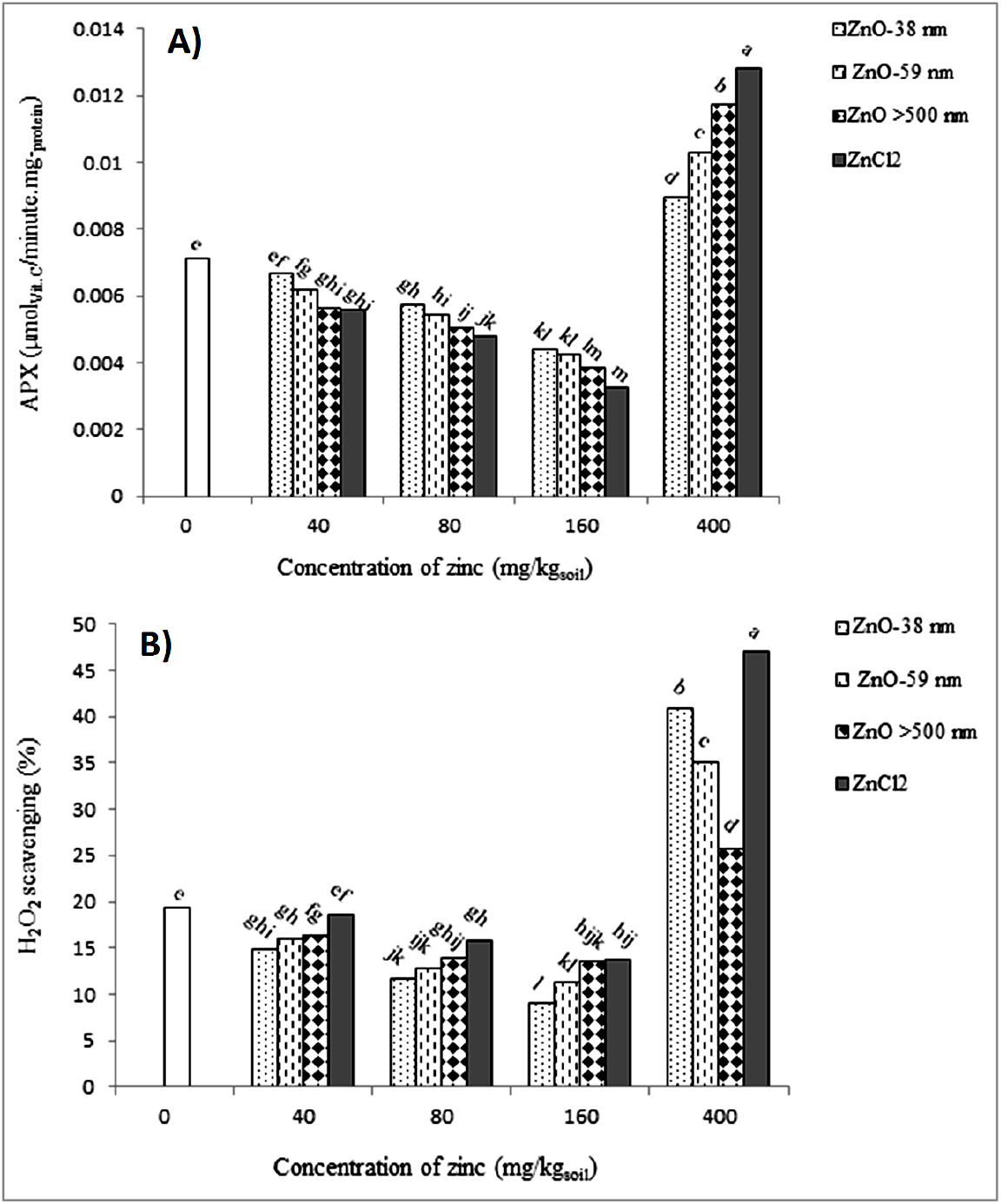
Changes in leaf ascorbate peroxidase (APX) (**A**), and H_2_O_2_ scavenging (**B**) activities in soil grown soybean treated with different types of zinc compounds (spherical ZnONP-38nm, florallike ZnONP-59nm, rod-like ZnONP>500nm, and Zn^2+^ ions) as a function of concentrations (0, 40, 80, 160, 400 mg Zn/kg soil). Error bars represent mean ± SD. Different letters above the bar indicate significant difference at *p*<0.05.

For the APX activity, which measured the ascorbate oxidation rate in the presence of added H_2_O_2_, we observed particle size-, morphology- and concentration-dependent influence of ZnONPs in soybean leaf (**Fig. 5A**). With increasing concentrations, APX activity was significantly reduced up to 160 mg/kg for all the Zn compounds tested; whereas at 400 mg/kg, APX activity increased significantly for all Zn compounds compared to control (**Fig. 5A**). Further, spherical 38nm ZnONPs were the most protective than all other Zn compounds tested (floral-like 59nm ZnONPs, rod-like >500nm ZnONPs, and Zn^2+^ ions) up to 160 mg/kg; but at the highest concentration (400 mg/kg) the toxicity trend reversed, i.e., Zn^2+^ ions were most protective than spherical 38nm ZnONPs. Overall, at 400 mg/kg all Zn compounds showed significantly higher APX activity (protective response) compared to control (**Fig. 5A**). Moreover, the APX responses for all Zn compounds were non-linear (**Fig. 5A;** see SI **Table S1** for R-squared values).

Our analysis of H_2_O_2_ scavenging ability of different ZnONPs showed particle size-, morphology- and concentration-dependent influence of ZnONPs in soybean leaf (**Fig. 5B**). With increasing concentrations, H_2_O_2_ scavenging was significantly reduced up to 160 mg/kg for all the Zn compounds tested; whereas at 400 mg/kg, the scavenging activity increased significantly for all Zn compounds compared to control (**Fig. 5B**). Further, spherical 38nm ZnONPs were least effective in scavenging leaf H_2_O_2_ than all other Zn compounds tested (floral-like 59nm ZnONPs, rod-like >500nm ZnONPs, and Zn^2+^ ions) up to 160 mg/kg; but at the highest concentration (400 mg/kg) the scavenging trend reversed for NPs, i.e., spherical 38nm ZnONPs were most effective in scavenging leaf H_2_O_2_ than two other ZnONPs tested (floral-like 59nm ZnONPs, rod-like >500nm ZnONPs). Whereas, at 400 mg/kg Zn^2+^ ions showed the most H_2_O_2_ scavenging potential than all three types of ZnONPs. Overall, leaf H_2_O_2_ scavenging responses for all Zn compounds were nonlinear (**Fig. 5B;** see SI **Table S1** for R-squared values).

## Discussion

Previous studies have documented variable toxicity of ZnONPs in soybean and other plants. However, most of these studies only focused on one size ZnONPs, thereby limiting our understanding about how some key physicochemical factors such as size, morphology (shape) and surface charge could play a role in nanotoxicity. For example, exposure to 18 nm ZnONPs led to increased chlorophyll content, shoot height, and grain yield in winter wheat, while plant biomass remained unaffected (Dimkpa et al., 2018). Foliar application of nanocomposites of ZnO, CuO, and B_2_O_3_ showed a significant increase in soybean biomass, grain count, grain biomass, and uptake of micro- and macro nutrients (Dimkpa et al., 2017a). Pod frequency increased in soybean exposed to ZnONPs, whereas the size and mean number of seed per pod did not differ between treatments (Priester et al., 2012). Wheat biomass and grain yield were promoted by 72% and 55%, respectively, upon ZnSO_4_ treatment, while ZnONPs treatment led to 63% and 56% increase in biomass and grain yield, respectively, compared to control (Du et al., 2019). However, in *Vigna unguiculata* grown in soil amended with soluble Zn^2+^ ions or ZnONPs showed no significant difference in plant growth between the two Zn compounds (Wang et al., 2013). Plant uptake and transport of NPs to aerial plant parts have been previously documented for other types of NPs exposures (Mueller and Nowack, 2008; Shang et al., 2019), leading to decreased plant biomass (Ma et al., 2015a; Shang et al., 2019). As previously discussed, Zn is required for plants to execute many physiological activities such as the biosynthesis of proteins and enzymes, chlorophyll, and normal functioning of the metabolic processes (Singh et al., 2018). In this study, seed yield was highest for spherical 38nm ZnONPs compared to floral-like 59nm ZnONPs or rodlike >500nm ZnONPs (**Fig. 2**) up to 160 mg Zn/kg treatments. Seed yield was generally reduced with Zn^2+^ ions treatments compared to all three types of ZnONPs treatments (**Fig. 2**), suggesting an inhibitory effect of Zn^2+^ ions on soybean seed development compared to ZnONPs evaluated. While we did not measure potential uptake of ZnONPs in soybean, it is possible that the biochemical processes discussed above (Chandra et al., 2014; Shaw and Hossain, 2013; Da Costa and Sharma, 2016) might have been altered with exposure to soluble Zn^2+^ ions and ZnONPs, potentially promoting (at 40-160 mg/kg) or inhibiting (at 400 mg/kg) seed yield. Taken together, our results suggest differential effects of ZnONPs compared to Zn^2+^ ions treatments (**Fig. 2**). Further, reduced energy demand due to diminished antioxidative enzymes (SOD, CAT, POX, and APX) (Kurutas, 2016; Yusefi-Tanha et al., 2020) being expressed up to 160 mg/kg of ZnONPs or Zn^2+^ ions treatments might explain increased seed yield, and the inverse response at 400 mg/kg of ZnONPs or Zn^2+^ ions treatment.

Tolerance of plants to metal-based nanoparticles exposure is determined by their ability to protect efficient antioxidant systems meant for scavenging or removal of the excess ROS (H_2_O_2_, OH^-^) within the cells and tissues. Previous studies have shown lower toxicity of bulk ZnO compared to ZnONPs (Amooaghaie et al., 2016; Mukherjee et al., 2014a), and that ZnONPs of small size were transformed into larger sized aggregates within the plant cell, complicating toxicity assessment (Lee et al., 2013). Toxicity induced by the higher concentration of Zn (400 mg Zn/kg) is manifested as potential interference with key antioxidative enzyme activities as depicted in **Figs. 2–5A**. H_2_O_2_ is toxic to the cell, particularly, at higher levels, so plants must maintain a balance in the cell to minimize its level to avoid oxidative stress and potential lipid peroxidation (Das and Roychoudhury, 2014). Previous works have shown increased ROS with exposure to higher concentrations of <100nm ZnONPs (500 mg Zn/kg) in wheat (Dimkpa et al., 2012) or 90nm ZnONPs (800 mg/kg soil) in maize (Liu et al., 2015). The hydroxyl (OH^-^) radical, the most reactive among all ROS, is produced from H_2_O_2_ as an outcome of dismutation, and is able to separate hydrogen atom from a methylene (CH2) group in the lateral chains of polyunsaturated fatty acids of membrane lipids thus initiating lipid peroxidation (Shaw et al., 2014). Our observation of particle size-, morphology- and concentration-dependent influence of ZnONPs on MDA accumulation in soybean as a response to H_2_O_2_ synthesis and lipid peroxidation in leaf is unique to nanophytotoxicity study (**Fig. 3**). Nair and Chung (2014a) observed a significant increase in H_2_O_2_ in soybean seedling roots upon exposure to different CuONP concentrations. Reduced lipid peroxidation (MDA level) observed with exposure to rod-like ZnONPs >500nm, especially at the highest concentration (400 mg/kg), may indicate the lack of particle transport in the plant owing to its larger particle size and rod-like morphology, which may have conferred significant protection to cell membrane integrity compared to other two types of ZnONPs and Zn^2+^ ions. These results are consistent with the previous study that showed a threshold for protective effect beyond which toxicity can manifest for ZnONPs in onion root cells (Kumari et al., 2011).

Hernandez-Viezcas et al. (2011) reported increased CAT activity in root, shoot and leaves at different concentrations (500-4000 mg/L) of 10nm ZnONPs treatments, but at concentrations higher than 500 mg/L APX activity was reduced. *Allium cepa* exposed to ZnONPs of~85nm (200, 400, and 800 mg/L) led to decreased CAT activity, but did not affect ROS level and glutathione peroxidase activity (Ghosh et al., 2016). In floating aquatic fern (*Salvinia natans*), exposure to 25nm ZnONPs (1-50 mg/L) showed no effect on plant growth, but led to increased SOD and CAT activities at the highest concentration tested (Hu et al., 2014), suggesting increased oxidative stress responses of fern at 50 mg Zn/L. At 500 and 750 mg Zn/kg, reduced CAT enzyme activity in stem and leaf were documented for different Zn compounds in alfalfa (Bandyopadhyay et al., 2015). Our findings showed that MDA and multiple antioxidant biomarkers (SOD, CAT, and POX) were altered in a similar fashion by the Zn compound type, concentrations, and their interactions (**Tables 2, 3; Fig. 3**). More specifically, the results showed particle size-, morphology-, and concentration-dependent influence of ZnONPs on MDA and antioxidant biomarkers, including SOD, CAT and POX, in soybean grown for 120 days. Overall, up to 160 mg Zn/kg treatments significant protective effects were observed in soybean by all Zn compound types tested.

**Table 3.** Analysis of variance (*p*-value) for antioxidant enzymes activity of soybean grown in soil treated with different concentration of zinc compound types.

| Source of variation | SOD | CAT | POX | APX |
| --- | --- | --- | --- | --- |
| Zinc compound type (Zn<sub>type</sub>) | <.0001 | <.0001 | <.0001 | ns |
| Concentration (C) | <.0001 | <.0001 | <.0001 | <.0001 |
| Zn<sub>type</sub> × C | <.0001 | <.0001 | <.0001 | <.0001 |

As a key antioxidant in the chloroplast, cytosol and mitochondria (Demidchik, 2015), SOD catalyzes the superoxide radical (O_2_^-^) and converts to H_2_O_2_ and O_2_, thus playing a major role in counteracting oxidative stress (Shekhawat, 2013; Ahmed et al., 2018). The results showed that increased leaf SOD activity (O_2_^-^ + 2H^+^ → H_2_O_2_ + O_2_) at 400 mg Zn/kg treatments might have promoted H_2_O_2_ production as shown in **Fig. 3A**. However, overall SOD activity was lower up to 160 mg Zn/kg treatments for all Zn compounds tested (**Fig. 3A**). Superoxide (O_2_^-^) levels were found to increase with 400-3,200 mg Zn/kg of ZnONPs (90±10 nm) and a significant increment in SOD activity was documented at the highest dose in maize (Wang et al., 2016a). In *Gossypium hirsutum,* increased activities of SOD and POX were reported with a subsequent decrease in lipid peroxidation with ZnONPs treatments (Venkatachalam et al., 2017). These results are consistent with our study. Enzymes such as CAT, POX and APX are known to catalytically reduce H_2_O_2_ into water and oxygen within the cell chloroplasts, cytosol, mitochondria, and/or peroxisomes (Anjum et al., 2016). These and other antioxidants may act independently or together via a crosstalk and reduce the extent of oxidative damage and toxicity in plants (Ahmed et al., 2017a and 2017b). CAT, POX, and APX enzymes seem to function concurrently. Herein, low APX activity (**Fig. 5A**) followed higher CAT and POX activities for ZnONPs and Zn^2+^ ions (**Figs. 4B,C**) - a finding consistent with our previous results documented for CuONPs treatments in soybean (Yusefi-Tanha et al., 2020). The overall trend of APX activity upon Zn compound treatments were reversed compared to MDA accumulation and SOD, CAT and POX responses in soybean leaf (**Fig. 5A**). Whereas, the scavenging of H_2_O_2_ by the Zn compounds showed a trend similar to MDA, SOD, CAT and POX responses (**Fig. 5B**). Further, we documented a clear threshold of 160 mg Zn/kg beyond which the responses of soybean upon exposure to all Zn compounds were found to be negative for all the endpoints measured. Our use of the high purity, same (triclinic) crystal structure and (negative) surface charge for all the three sizes and morphologically different ZnONPs enabled us to specifically test for the size- and morphology-dependent effects of ZnONPs in soybean. Our results showed a significant particle size, morphology and concentration dependent influence of ZnONPs on seed yield, lipid peroxidation, and various antioxidant biomarkers in soil-grown soybean. The findings further suggest unique toxicity of ZnONPs compared to ionic Zn^2+^ in soybean.

An estimated 50% of arable soil being used in agriculture may contain reduced amount of soluble Zn, leading to reduced crop yield and poor nutritional quality of grains and derivatives (Moreira et al., 2018). Organic matter is known to influence soil Zn availability to plants with ZnONPs or bulk Zn treatments (Moghaddasi et al., 2017; Medina-Velo et al., 2017; Dimpka et al., 2020). Recently, Dimkpa et al. (2020) documented beneficial effects of 18nm ZnONPs on various endpoints (shoot length, plant height, shoot biomass, chlorophyll, grain yield, shoot/grain Zn^2+^ uptake, grain Ca^2+^ and Mg^2+^ uptake) measured in wheat under simulated drought (at 40% field moisture capacity) and/or with organic manure (cow dung) amendment. At half the concentration, ZnONPs (2.17 mg ZnO/kg) showed similar or greater positive effects than bulk ZnO treatments (>1000nm powder). These findings are significant in that potentially higher reactivity of ZnONPs may prove more sustainable (on a mass-by-mass basis) compared to bulk Zn fertilization of soil (Dimkpa et al., 2020). Our findings also suggest the prospect of ZnONPs to be used as a nanofertilizer (up to 160 mg Zn/kg) for supplementing Zn element to plants (Singh et al., 2013), reducing oxidative stress, and promoting seed yield. Subsequently, through edible plant parts such as leaves and seeds, Zn is bioavailable to humans at as high as 70% while average absorption is about 33% (Turnlund et al., 1984; Cousins, 1985; FAO/WHO, 2004). As an essential micronutrient, Zn deficiency has been linked to hypogonadism, anemia, and stunted growth in humans (Roohani et al., 2013); thus, nano-fortification of plants during growth with the required quantity of Zn (this may depend on soil type) could be a viable concept—one that might be a game changer, particularly, for people who rely on high plant-based diets such as cereals and beans and those who consume less meat (Roohani et al., 2013).

Assessment of the R-squared values for all the endpoints measured showed nonlinear concentration-response curves (R^2^<0.65) (see SI **Table S1**). Further, our results demonstrated no observed adverse effect level (NOAEL) of 160 mg/kg for all three different ZnONPs types, including for Zn^2+^ ions, for all the endpoints measured (see TOC art), except for H_2_O_2_ scavenging (**Fig. 5B**). The protective effects conferred by ZnONPs up to 160 mg/kg in soybean can be attributed to the role of Zn as an essential micronutrient in many enzymes and proteins, which are key for plant growth, development and crop yield (Moreira et al., 2018).

## Conclusion

In summary, our findings demonstrated a significant particle size-, morphology- and concentration-dependent influence of ZnONPs on seed yield, lipid peroxidation, and various antioxidant biomarkers measured in soil-grown soybean. The spherical 38nm ZnONPs were the most protective compared to the floral-like 59nm ZnONPs, rod-like >500nm ZnONPs, or Zn^2+^ ions, particularly up to 160 mg/kg. However, at the highest concentration of 400 mg/kg, spherical 38nm ZnONPs elicited the highest oxidative stress responses (H_2_O_2_ synthesis, MDA, SOD, CAT, POX) in soybean compared to the other two morphologically different ZnONPs tested. The concentration-response curves for the three types of ZnONPs and Zn^2+^ ions were nonlinear for all the endpoints evaluated. Our results also suggest differential nano-specific toxicity of ZnONPs compared to ionic Zn^2+^ toxicity in soybean. Furthermore, the higher NOAEL value of 160 mg/kg indicates the potential for using ZnONPs as a nanofertilizer for crops grown in Zn deficient soils to improve crop yield, food quality and address malnutrition, globally.

## Acknowledgements

This study was conducted at the Department of Agronomy, Shahrekord University, Iran. The authors would like to thank Shahrekord University for providing financial support. LRP gratefully acknowledges funding support from East Carolina University (grant # 111101 to LRP).

## Supplementary Information

**Table S1.** R-squared values based on the linear regression lines for multiple parameters tested for different sized ZnONPs and ZnCl_2_ treatments in soybean. To determine if the concentration-response curves were linear (monotonic) or nonlinear (nonmonotonic), we coupled visual inspection of the curves with a simple decision rule: if the co-efficient of determination (R-squared) value for the linear regression line is 65% or higher the concentration-response curves were deemed linear, suggesting that the plant response changes linearly with the concentration applied. ‘L’ denotes linear, and ‘NL’ denotes nonlinear, concentration-response relationships.

| Parameters | R-squared |  |  |  |
| --- | --- | --- | --- | --- |
|  | ZnONP-38 nm | ZnONP-59 nm | ZnONP>500 nm | ZnCl<sub>2</sub> |
| **Seed production** | 0.22 (NL) | 0.20 (NL) | 0.066 (NL) | 0.32 (NL) |
| **H<sub>2</sub>O<sub>2</sub>** | 0.46 (NL) | 0.33 (NL) | 0.044 (NL) | 0.63 (NL) |
| **MDA** | 0.50 (NL) | 0.42 (NL) | 0.047 (NL) | 0.64 (NL) |
| **SOD** | 0.36 (NL) | 0.22 (NL) | 0.036 (NL) | 0.61 (NL) |
| **CAT** | 0.29 (NL) | 0.21 (NL) | 0.10 (NL) | 0.45 (NL) |
| **POX** | 0.37 (NL) | 0.34 (NL) | 0.26 (NL) | 0.51 (NL) |
| **APX** | 0.16 (NL) | 0.33 (NL) | 0.42 (NL) | 0.42 (NL) |
| **H<sub>2</sub>O<sub>2</sub> Scavenging** | 0.51 (NL) | 0.49 (NL) | 0.37 (NL) | 0.64 (NL) |

